# Aberrant Visual Population Receptive Fields in Human Albinism

**DOI:** 10.1101/572289

**Authors:** Ethan J. Duwell, Erica N. Woertz, Jedidiah Mathis, Joseph Carroll, Edgar A. DeYoe

## Abstract

Retinotopic organization is a fundamental feature of visual cortex thought to play a vital role in encoding spatial information. One important aspect of normal retinotopy is the representation of the right and left hemifields in contralateral visual cortex. However, in human albinism, many temporal retinal afferents decussate pathologically at the optic chiasm resulting in partially superimposed representations of opposite hemifields in each hemisphere of visual cortex. Previous fMRI studies in human albinism suggest that the right and left hemifield representations are superimposed in a mirror-symmetric manner. This should produce imaging voxels which respond to two separate regions in visual space mirrored across the vertical meridian. However, it is not yet clear how retino-cortical miswiring in albinism manifests at the level of single voxel population receptive fields. Here we used fMRI retinotopic mapping in conjunction with population receptive field (pRF) modeling to fit both single and dual pRF models to the visual responses of voxels in visual areas V1-V3 of five subjects with albinism. We found that subjects with albinism (but not controls) have sizable clusters of voxels with dual pRFs consistently corresponding to, but not fully coextensive with regions of hemifield overlap. These dual pRFs were typically positioned at roughly mirror image locations across the vertical meridian but were uniquely clustered within the visual field for each subject. We also found that single pRFs are larger in albinism than controls, and that single pRF sizes in the central visual field were anti-correlated with subjects’ foveal cone densities. Finally, dual pRF and aberrant hemifield representation characteristics varied significantly across subjects with albinism suggesting more central heterogeneity than previously appreciated.

## INTRODUCTION

Retinotopic organization is one of the most fundamental and well-described organizational principles of visual cortex. Present at every level of the visual hierarchy, the retinotopic organization of visual cortex is thought to play a vital role in the brain’s ability to encode visual-spatial information. The establishment of retinotopic organization requires a surprising degree of connectional specificity during development. One promising route toward better understanding the development and functional organization of retinotopic maps is to study pathological syndromes where normal retinotopy is disrupted. In these cases, we can quantify exactly how retinotopic organization differs from normal and thereby provide a physiological basis for understanding and predicting potentially aberrant perceptual consequences.

Albinism is a well-known genetic syndrome characterized by disrupted melanin synthesis which causes hypopigmentation of the eyes, and often the skin and hair. In addition, albinism is also associated with aberrant decussation of the temporal retinal afferents at the optic chiasm such that each cortical hemisphere receives substantial input from both the right and left visual hemifields of the same eye (Hoffmann, Tolhurst, Moore, & Morland, 2003; Kaule, Wolynski, Gottlob, Stadler, Speck, Kanowski, Meltendorf, Behrens-Baumann, & Hoffmann, 2014; Morland, Baseler, Hoffmann, Sharpe, & Wandell, 2001). This results in overlaid retinotopic maps of significant portions of *opposite* hemifields within each hemisphere of occipital visual cortex. This differs from the arrangement in healthy, control subjects, where each hemisphere contains superimposed maps of the *same* (contralateral) hemifield from each eye (Horton & Hoyt, 1991). Thus, the pattern of miswiring in albinism represents a major disruption of normal retinotopic organization.

Previous fMRI mapping studies in albinism suggest that the right and left hemifield representations are superimposed in a precise manner such that each voxel responds to two regions at roughly mirror-image positions across the vertical meridian (Hoffmann et al., 2003). These results are generally consistent with the “true albino” pattern of hemifield superposition previously described in albino monkeys in which ocular dominance columns are supplanted by hemifield dominance columns (Guillery, Hickey, Kaas, Felleman, Debruyn, & Sparks, 1984; Hoffmann & Dumoulin, 2015). This columnar pattern was recently confirmed in high-resolution fMRI studies in human achiasma (Olman, Bao, Engel, Grant, Purington, Qiu, Schallmo, & Tjan, 2018) but has yet to be demonstrated in albinism. More recently, investigators have used single voxel population receptive field (pRF) modeling techniques to demonstrate the existence of single voxels with mirror symmetric bilateral (dual) pRFs in achiasma and FHONDA syndrome (Ahmadi, Fracasso, van Dijk, Kruijt, van Genderen, Dumoulin, & Hoffmann, 2019; Hoffmann, Kaule, Levin, Masuda, Kumar, Gottlob, Horiguchi, Dougherty, Stadler, Wolynski, Speck, Kanowski, Liao, Wandell, & Dumoulin, 2012). These studies have assumed that subjects with albinism should present with the same mirror symmetric, dual pRF topographies. However, this may not be the case as the genetic determinants and nature of retinocortical miswiring differ in achiasma, FHONDA, and albinism (Ahmadi et al., 2019; Hoffmann & Dumoulin, 2015), and recent reports conflict regarding the existence of dual pRFs in albinism (Ahmadi, Herbik, Wagner, Kanowski, Thieme, & Hoffmann, 2019; Alvarez, Smittenaar, Handley, Liasis, Sereno, Schwarzkopf, & Clark, 2019). Moreover, albinism itself is a heterogeneous syndrome with a wide variety of genetic and phenotypic subtypes (Montoliu, Grønskov, Wei, Martinez-Garcia, Fernandez, Arveiler, Morice-Picard, Riazuddin, Suzuki, Ahmed, Rosenberg, & Li, 2014; Oetting, Summers, & King, 1994; Prieur & Rebsam, 2017; Simeonov, Wang, Wang, Sergeev, Dolinska, Bower, Fischer, Winer, Dubrovsky, Balog, Huizing, Hart, Zein, Gahl, Brooks, & Adams, 2013; Wilk, McAllister, Cooper, Dubis, Patitucci, Summerfelt, Anderson, Stepien, Costakos, Connor, Wirostko, Chiang, Dubra, Curcio, Brilliant, Summers, & Carroll, 2014). In a previous study (Woertz, Wilk, Duwell, Mathis, Carroll, & DeYoe, 2020), we described significant variation in the pattern of hemifield eccentricity mapping in subjects with albinism, thus suggesting that pRF properties in albinism may vary considerably across subjects.

Albinism is also associated with reduced foveal cone density, foveal hypoplasia, and low visual acuity(Wilk et al., 2014; Wilk, Wilk, Langlo, Cooper, & Carroll, 2017; Woertz et al., 2020). Lower cone density in the foveal center results in coarser sampling of visual space and correspondingly low central visual acuity (Rossi & Roorda, 2010). This coarser sampling in the fovea may manifest cortically in the form of larger visual pRFs. In this way pRF modeling may inform our understanding of the cortical substrates of visual acuity deficits in the albinism population.

In this study, we used traditional fMRI retinotopic mapping in combination with population receptive field modeling of voxels in V1-V3 of subjects with genetically confirmed albinism. We tested the hypothesis that cortical retinotopic organization in albinism is aberrant and that this is associated with voxels in visual cortex having bilateral (dual) pRFs. Our results reveal imaging voxels with dual pRFs in albinism consistently associated with the superimposed opposite hemifield representations and responding to fields at *roughly* symmetrical locations across the vertical meridian. However, not all voxels within the hemifield overlap zones had dual receptive fields. Some were adequately fit by a single pRF model. Finally, our results also showed that single pRFs are consistently larger in albinism than controls, and that single pRF sizes in the central visual field are highly anti-correlated with subjects’ foveal cone densities.

## METHODS

### Subjects

The subject cohort and data used in this study are the same as in our recent study of cortical magnification in albinism (Woertz et al., 2020). Six subjects with albinism (4 females, 2 males; aged 15-31 years) with minimal nystagmus and five control subjects with no prior ocular or cortical pathology (2 females, 3 males; aged 20-25 years) were recruited for this experiment. One subject with albinism was excluded from further analysis due to significant motion artefacts in the fMRI data (male, age 15 years). Genetic information and demographics for each albinism subject are listed in **Table 1** below, and data characterizing fixation stability in each subject are presented in **Table 2**. Retinal features and cortical magnification in these subjects are described in detail elsewhere as are the methods used to determine best corrected visual acuity (BCVA), albinism subtype, and genetic mutations (Wilk et al., 2014; Wilk et al., 2017; Woertz et al., 2020). It is also notable that subjects 2 and 4 are siblings with identical mutations. The study was in accordance with the Declaration of Helsinki and approved by the Institutional Review Board of the Medical College of Wisconsin. All subjects provided written informed consent after explanation of the nature and possible risks and benefits of the study.

**Table 1:**
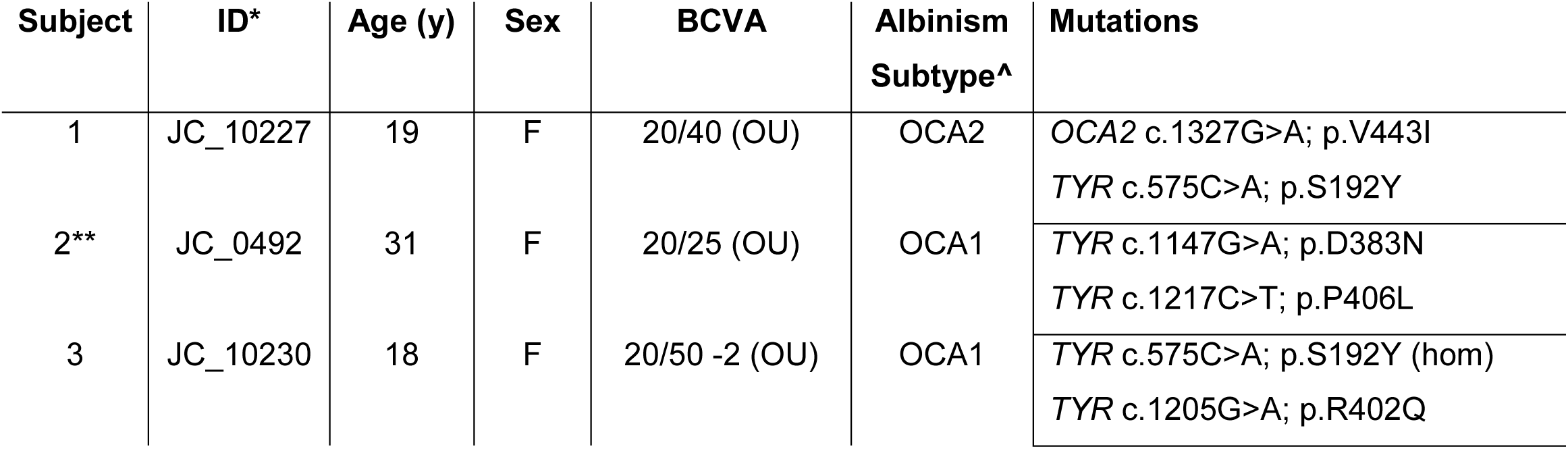
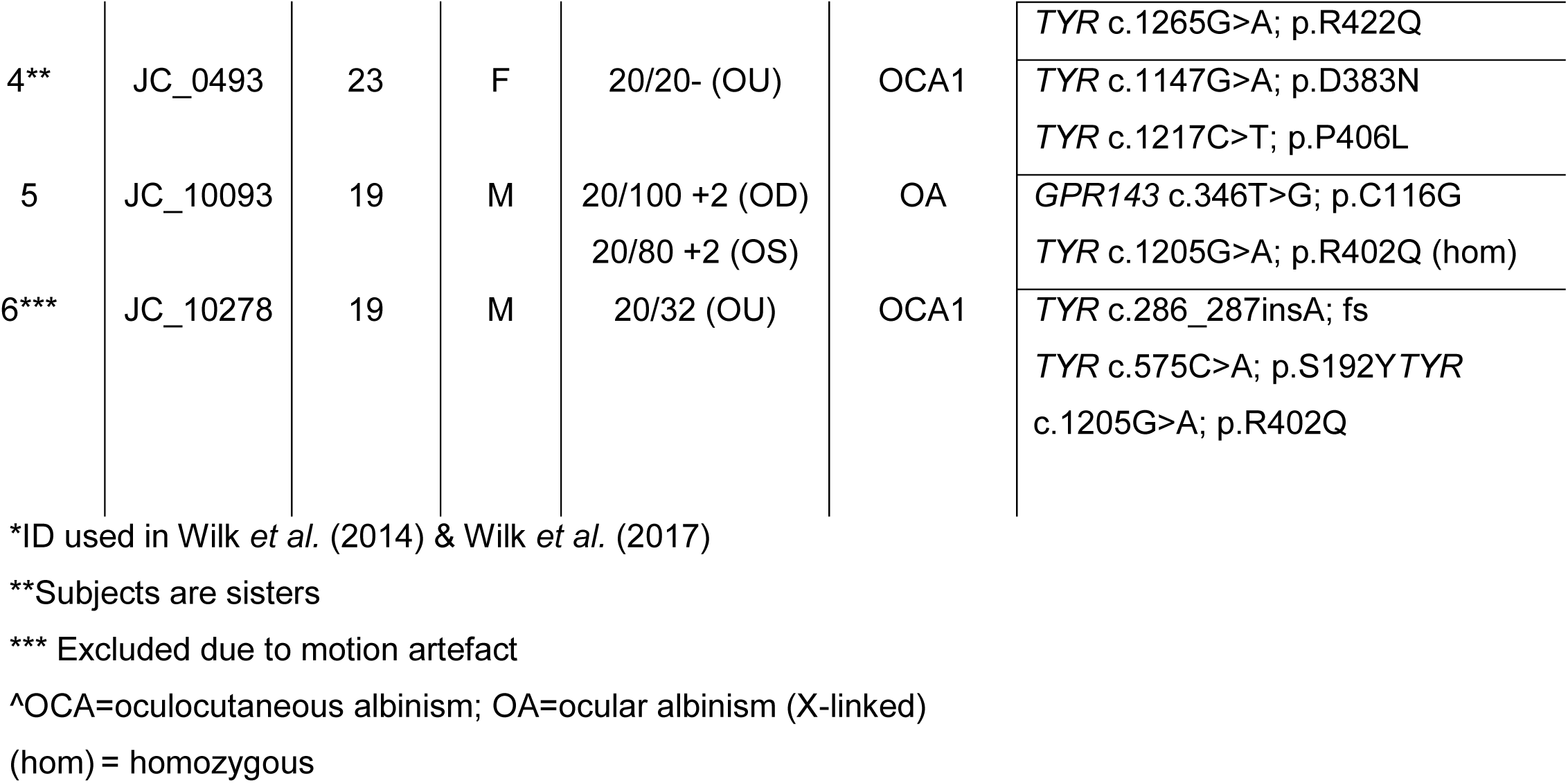
Genetics and Demographics for Subjects with Albinism

**Table 2:** Fixational stability in subjects with albinism

| Subject | ID | OD |  | OS |  |
| --- | --- | --- | --- | --- | --- |
|  |  | 50% BCEA | 95% BCEA | 50% BCEA | 95% BCEA |
| 1 | JC_10227 | 0.24 | 1.05 | 0.45 | 1.95 |
| 2 | JC_0492 | 0.04 | 0.19 | 0.06 | 0.28 |
| 3 | JC_10230 | 1.92 | 8.35 | 0.62 | 2.68 |
| 4 | JC_0493 | 0.84 | 3.66 | 0.14 | 0.597 |
| 5 | JC_10093 | 3.68 | 16.01 | 5.97 | 25.95 |

OD = right eye; OS = left eye; BCEA = bivariate contour ellipse area. All values are expressed in degrees<sup>2</sup>.

### Fixation Testing

An in-depth description of the fixation testing method performed on these subjects is provided in (Woertz et al., 2020). In brief, subjects fixated on a small white cross while their fixation stability was monitored using the fixation test module of an OPKO combined scanning laser ophthalmoscope (SLO). These retinal imaging data were then used to compute the 50% and 95% bivariate contour ellipse areas (BCEA) that characterize each subject’s fixation stability.

### fMRI Visual Stimuli

Note: The following descriptions of the fMRI visual stimuli, fMRI stimulus paradigm, fMRI acquisition, phase encoded retinotopic maps, and visual area mapping are nearly identical to those described in our recent study of cortical magnification in albinism (Woertz et al., 2020). This study utilizes the same fMRI data.

All visual stimuli for fMRI were presented on a back-projection screen mounted on the MRI head coil. A BrainLogics BLMRDP-A05 MR digital projector was used with a ViSaGe MKII visual stimulus generator (Cambridge Research Instruments) in conjunction with custom MATLAB software. Stimuli subtending a maximum of 20° eccentricity included conventional expanding ring and rotating wedge retinotopic mapping stimuli (DeYoe, Carman, Bandettini, Glickman, Wieser, Cox, Miller, & Neitz, 1996). Rings and wedges were composed of black and white counterphase flickering (8 Hz) circular checkerboards with check size and ring width scaled with eccentricity. Stimuli were photopic and presented on a uniform gray background. All subjects were instructed to continually fixate a marker at the center of the screen. To enhance stable fixation, thin, black radial lines extending from fixation to the edge of the display were present continuously during all tasks.

To avoid collecting redundant MRI data on control subjects, we used previously acquired retinotopic mapping data even though it was obtained with a slightly different experimental protocol than for subjects with albinism. However, due to the temporal phase mapping methods used in this study, retinotopic parameters (eccentricity, polar angle) that are encoded by the timing of the fMRI responses did not appear to be significantly affected by the stimulus differences. For subjects with albinism, the wedge stimuli subtended 45° polar angle, whereas for control subjects the wedges subtended 90°. All subjects viewed both ring and wedge stimuli binocularly with full-field stimulation. Additionally, for subjects with albinism the expanding ring stimuli were presented to the right and left hemifields in separate runs and were tested separately for each eye. The hemifield ring stimuli were identical to the full field version, except that one hemifield was masked to match the grey background. For control subjects, the ring stimulus expanded from the center to the periphery in 40 s. and was repeated five times per run. For subjects with albinism, both the full-field and hemifield ring stimuli expanded from 0.8° eccentricity to the periphery in 60 s. and were repeated five times per run. For subjects with albinism, the center of the display consisted of a circular black and white disc (similar to a radioactivity symbol) with a radius of 0.8° that flickered at random intervals not synchronized to the rings or wedges presentation. To control attention, subjects were instructed to press and hold a button whenever the central disc flickered.

### fMRI Stimulus Paradigm

Control subjects completed all imaging during a single session. Subjects with albinism completed imaging during two sessions: the right eye hemifield expanding ring tasks in the first session, and all remaining tasks in the second session. All monocular hemifield runs were repeated five times and binocular full-field runs were repeated three times. For monocular stimuli, repetitions of the right and left hemifield stimuli were interleaved; for full-field stimuli, repetitions of the expanding ring and rotating wedge were interleaved. After each fMRI run, the subject was asked to rate their alertness on a scale from 1-5 (1 being asleep and 5 being fully awake). This measure was used to control for subjects’ alertness which can affect the quality of data. We used this measure as an exclusion criterion. However, no subjects were excluded from this study for poor alertness.

### fMRI Acquisition

MRI scans were obtained at the Medical College of Wisconsin using a 3.0 Tesla General Electric Signa Excite 750 MRI system equipped with a custom 32-channel RF/gradient head coil. BOLD fMRI images were acquired with a T2*-weighted gradient-echo EPI pulse sequence (TE = 25 ms, TR = 2 s, FA = 77°). The 96 x 96 acquisition matrix (Fourier interpolated to 128 × 128) had frequency encoding in the right-left axial plane, phase encoding in anterior-posterior direction and slice selection in the axial direction. The FOV was 240 mm and included 29 axial slices covering the occipital lobe and adjacent portions of the temporal and parietal lobes with a slice thickness of 2.5 mm, yielding a raw voxel size of 1.875 × 1.875 × 2.5 mm. For anatomical scans, a T1-weighted, spoiled gradient recalled at steady state (SPGR), echo-planar pulse sequence was used (TE = 3.2 ms, TR = 8.2 ms, FA = 12°) with a 256 × 224 acquisition matrix (Fourier interpolated to 256 x 256). The FOV was 240 mm, and 180 slices with a slice thickness of 1.0 mm, yielding voxel sizes of 0.938 × 0.938 × 1.0 mm^3^. The SPGR scans were subsequently resampled to 1.0 mm^3^. A sync pulse from the scanner at the beginning of each run triggered the onset of visual stimuli.

### Analysis Software

All fMRI data were analyzed using the AFNI/SUMA software package (version: AFNI_19.1.11, https://afni.nimh.nih.gov/) (Cox, 1996). Surface models were produced from the high resolution SPGR images using the ‘recon-all’ function in Freesurfer (version 5.1.0 and 5.3.0, http://surfer.nmr.mgh.harvard.edu/). Single voxel time-course modeling was performed using custom software in MATLAB (R2017b).

### fMRI Pre-processing

fMRI pre-processing was performed using a script generated by AFNI’s afni_proc.py command, and occurred in the following order: reconstruction, removal of before and after periods, slice timing correction, alignment and volume registration, smoothing, scaling, regression of motion parameters and linear trends, averaging. Before and after periods were removed using AFNI 3dTcat, and slice time shift correction was then performed using AFNI 3dTshift. To reduce alignment bias for scans acquired during either of the two imaging sessions in our albinism subjects, the reference SPGR anatomical images from both sessions were skull-stripped using AFNI 3dSkullStrip, aligned using AFNI align_epi_anat.py, and averaged using AFNI 3dMean to create an average reference anatomy for the two sessions. For control subjects, all data were acquired in a single session, so the creation of an average reference anatomy was not necessary. The alignment of functional runs for control subjects was otherwise identical to subjects with albinism. Rigid body alignment and volume registration were performed using AFNIs align_epi_anat.py and 3dVolreg respectively. During volume registration, volumes in each run were registered to the volume in that series having the least motion as computed by AFNI 3dToutcount. This volume was also used as the base EPI for aligning each functional run to the anatomical scan. The alignment and volume registration transformations were computed separately, concatenated, and then applied together. Registration parameters from 3dVolreg were used to compute motion magnitude time series to serve as motion regressors later in the pipeline. Each run was then smoothed with a 3.75 mm (full width at half max) Gaussian kernel using AFNI 3dMerge, and then brain-masked using 3dAutomask. After masking, the timeseries data were then scaled to range from 0-200 with a mean of 100. Linear trends were then removed from the scaled data and a regression analysis of the motion regressors was then performed using 3dDeconvolve. The resulting motion regression matrix was projected out of the time series data using 3dTproject. Finally, individual runs from each task were averaged using 3dMean.

### Phase Encoded Retinotopic Maps

The spatial distributions of significant fMRI responses for each functional task were displayed as phase encoded retinotopic activation maps. Significant responses were identified by cross correlating the empirical time course data for each voxel with a reference waveform using AFNI 3ddelay (Bandettini, Jesmanowicz, Wong, & Hyde, 1993; Datta & DeYoe, 2009; Saad, DeYoe, & Ropella, 2003). This analysis produces the correlation coefficient and phase values at the phase offset of maximum correlation for each voxel. The reference waveform used for this phase mapping procedure was a binary square wave describing the stimulus cycles convolved temporally with the “Cox Wide” estimation of the hemodynamic response function (HRF).

### Visual Area Mapping

The correlation analysis described above was performed on the smoothed, full field, phase encoded retinotopy data, and the results were projected onto cortical surface models in AFNI/SUMA. These phase encoded polar angle and eccentricity maps were used to identify and map visual areas V1-V3 in both albinism subjects and controls using criteria previously reported by a number of labs (Amano, Wandell, & Dumoulin, 2009; Arcaro, McMains, Singer, & Kastner, 2009; DeYoe et al., 1996; Engel, Glover, & Wandell, 1997; Hansen, Kay, & Gallant, 2007; Pitzalis, Galletti, Huang, Patria, Committeri, Galati, Fattori, & Sereno, 2006; Pitzalis, Sereno, Committeri, Fattori, Galati, Patria, & Galletti, 2010; Sereno, Dale, Reppas, Kwong, Belliveau, Brady, Rosen, & Tootell, 1995; Sereno, Pitzalis, & Martinez, 2001; Silver & Kastner, 2009; Swisher, Halko, Merabet, McMains, & Somers, 2007; Wandell, Dumoulin, & Brewer, 2007; Wandell & Winawer, 2011). Visual area ROIs drawn on the cortical surface models were then transformed back to the volumetric domain using AFNI’s 3dSurf2Vol.

### Identifying Cortical Zones of Right-Left Hemifield Overlap in Albinism

To identify cortical regions responding to both the right and left hemifields for each individual subject with albinism, the monocular right and left hemifield eccentricity maps from both eyes were thresholded to a minimum correlation coefficient of 0.45, binarized, and then logically combined using AFNI’s 3dcalc to generate hemifield overlap maps. Any voxel which responded above threshold in both a left and right hemifield stimulation condition was included in the hemifield overlap ROI. Voxels which responded to only a single hemifield were included in the non-overlap ROI.

### Population Receptive Field Modeling

Both single and dual Gaussian population receptive field (pRF) models were optimized for each responsive voxel to fit the full field rotating wedge time-course data in visual areas V1-V3. The procedure was inspired by methods developed by Dumoulin et al. (Dumoulin & Wandell, 2008) but was implemented independently using custom Matlab software (Puckett & DeYoe, 2015). The modeling algorithm uses properties of the stimuli, tasks, and BOLD hemodynamics in conjunction with estimates of the pRF to generate a predicted fMRI waveform. This predicted waveform was then fit to the empirical time-course data yielding an error signal which was used to drive an iterative optimization of the Gaussian model parameters so as to fit the empirical data most accurately. Single pRFs were modeled as a simple 2D Gaussian:

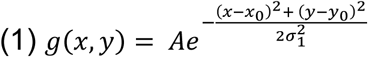

where (x_0,_ y_0_) is the center position, σ is the standard deviation of the distribution, and A is an amplitude scaling factor. Dual pRFs were modeled as the sum of two independent 2D Gaussians:

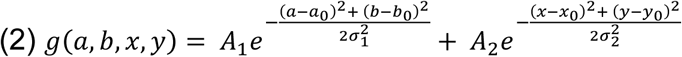

where (a_0_,b_0_) and (x_0_,y_0_) are the centers of the two Gaussians, σ_1_ and σ_2_ are the standard deviations of the two Gaussians, and A_1_ and A_2_ are the amplitude scaling factors.

The modeling procedure began by multiplying an initial pRF estimate with the spatial pattern of the stimulus and integrating over space at each time point to generate an ideal ‘neural’ response waveform. This ideal response was then convolved temporally with an estimate of the HRF to produce a predicted fMRI response. The residual sum squared error (RSS) between this predicted response and the empirical fMRI waveform was then computed. The RSS error signal drove an iterative optimization of the location, standard deviation, and amplitude parameters of the model searching for the combination which produced a minimum error in the fit between the model’s predicted timecourse and the voxel’s empirical timecourse. Optimization began with a coarse grid search of all possible (a_0_,b_0_) and (x_0_,y_0_) starting positions limited to the voxel’s preferred eccentricity as determined by the full field expanding ring experiment. Using the starting positions established by the coarse search, polar angle, σ, and amplitude (sensitivity) were refined in a two-pass, coarse-to-fine fashion using an unconstrained nonlinear optimization algorithm (Matlab’s fminsearch) to obtain an optimal fit to the empirical rotating wedge fMRI waveform.

This process was performed twice for each voxel: once with the single Gaussian pRF model, and again with the dual Gaussian model. We emphasize that for the dual Gaussian model, each Gaussian was optimized independently. Previous papers performing dual receptive field modeling in albinism, achiasma and other misrouting disorders have restricted the two receptive fields to be at mirror image locations either across the vertical meridian, horizontal meridian, or fixation (Ahmadi et al., 2019; Ahmadi et al., 2019; Alvarez et al., 2019; Hoffmann et al., 2012). By allowing the two Gaussians to be optimized independently, our model tested the possibility that dual pRFs in albinism may be arranged at any combination of angular positions at a given eccentricity.

### Voxel Selection

All of our quantitative analyses were performed only on voxels falling within our V1-V3 visual area ROIs. Selection criteria varied slightly in each separate analysis. To be included in our hemifield overlap analysis, voxel responses were required to achieve a correlation coefficient greater than 0.45 on at least 1 of the 4 monocular hemifield stimulation conditions. For voxels to be included in our pRF analysis, one of the two pRF models must have explained a minimum of 20% of the total timecourse variance for that voxel. The percent variance explained by each model was calculated using **Equation 4** below. For inclusion in our analysis of the incidence of dual pRF voxels in hemifield overlap vs. non overlap zones in albinism, voxels were required to meet both of the criteria above. In addition, as wedge and ring stimuli for the albinism group did not overlap the central 0.8 degrees of visual space (obscured by central fixation task stimulus), we excluded voxels responding to the central most 1 degree of visual space in both the albinism and control groups. This also served to eliminate artifacts known to occur at the center of gaze when using ring and wedge stimuli.

### Model Comparison

The fits of the single and dual pRF models for each voxel were compared using a partial *F*-test to determine whether the additional Gaussian parameters optimized in the dual pRF model resulted in a significant increase in variance explained. For each voxel, a partial *F*-static was computed using the following equation:

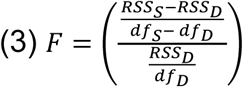

Where RSS_S_ is the *RSS* error from fitting the single pRF model, RSS_D_ is the *RSS* error from fitting the dual pRF model, *df*_*S*_ is the number of degrees of freedom in the single pRF model, and df_D_ is the number of degrees of freedom in the dual pRF model. *Df*_*S*_ and *df*_*D*_ included the number of free parameters optimized in the single and dual pRF models respectively plus the number of degrees of freedom used in the baseline motion/linear trends regression model projected out of the time series data during pre-processing. Voxels were considered to have dual pRFs if this partial *F*-statistic fell above the 99% confidence interval in the appropriate *F*-distribution. Voxels were considered to have single pRFs if their *F*-statistic fell below the 1% confidence interval. The percent variance explained (*%VE*) by each respective model was also computed on a voxel-wise basis using the following equation:

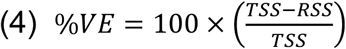

Where *TSS* is the total sum of squares for the voxel timecourse and *RSS* is the RSS error from the model fitting. Finally, the difference in variance explained (*ΔVE*) by the dual and single pRF models was computed for each voxel in the following manner:

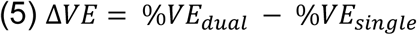

Where *%VE*_*dual*_ and *%VE*_*single*_ are the percent variance explained by the dual and single pRF models respectively.

### Dual pRF Symmetry

For all dual pRF voxels, the degree to which the dual pRFs were symmetric across the vertical meridian was assessed by computing the angle (*θ)* between the two pRF centers with respect to the horizontal meridian:

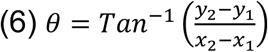

Where (*x*_*1*_, *y*_*1*_) are the Cartesian coordinates for pRF center 1 and (*x*_*2*_, *y*_*2*_) are the coordinates for pRF center 2. The absolute values of these angles can range from 0°-90°. Since the dual pRF components were constrained to be at the same eccentricity, an angle of 0°indicated symmetry across the vertical meridian, and 90°indicated symmetry across the horizontal meridian. Histograms of dual pRF angles were generated for each subject.

## RESULTS

### Fixational Stability

A detailed discussion of the fixation stability results in these subjects is provided in our recent paper on cortical magnification in albinism (Woertz et al., 2020). The 50% and 95% BCEA for each eye in each subject with albinism are shown in **Table 2**. All but one subject’s (subject 5) 50% BCEAs were below 3.14 deg^2^, the threshold previously used to define normal steady fixation (Fujii, De Juan, Humayun, Sunness, Chang, & Rossi, 2003; Woertz et al., 2020).

### Aberrant Retinotopic Organization in Albinism

As reported in the previous literature (Hoffmann et al., 2003; Kaule et al., 2014; Morland et al., 2001; Prieur & Rebsam, 2017), we observed overlapping representations of both left and right hemifields within each cerebral hemisphere of all our subjects with albinism. **Figure 1A,B** displays inflated cortical surface maps of the left occipital lobe in one representative subject with albinism (subject 5). In this case, right (A) and left (B) hemifield expanding ring stimuli were presented monocularly to the right eye. The eccentricity color code is shown in the upper left of **Figure 1A,B**. By visual inspection, the right and left hemifield eccentricity maps appeared to be grossly in register but the aberrant left hemifield representation (B) was less complete than the normal right hemifield representation (A) such that the two maps only partially overlap. The foveal representation in this subject, which should be at the occipital pole, was reduced relative to our control subjects, consistent with previous observations (Woertz et al., 2020).

**Figure 1:**
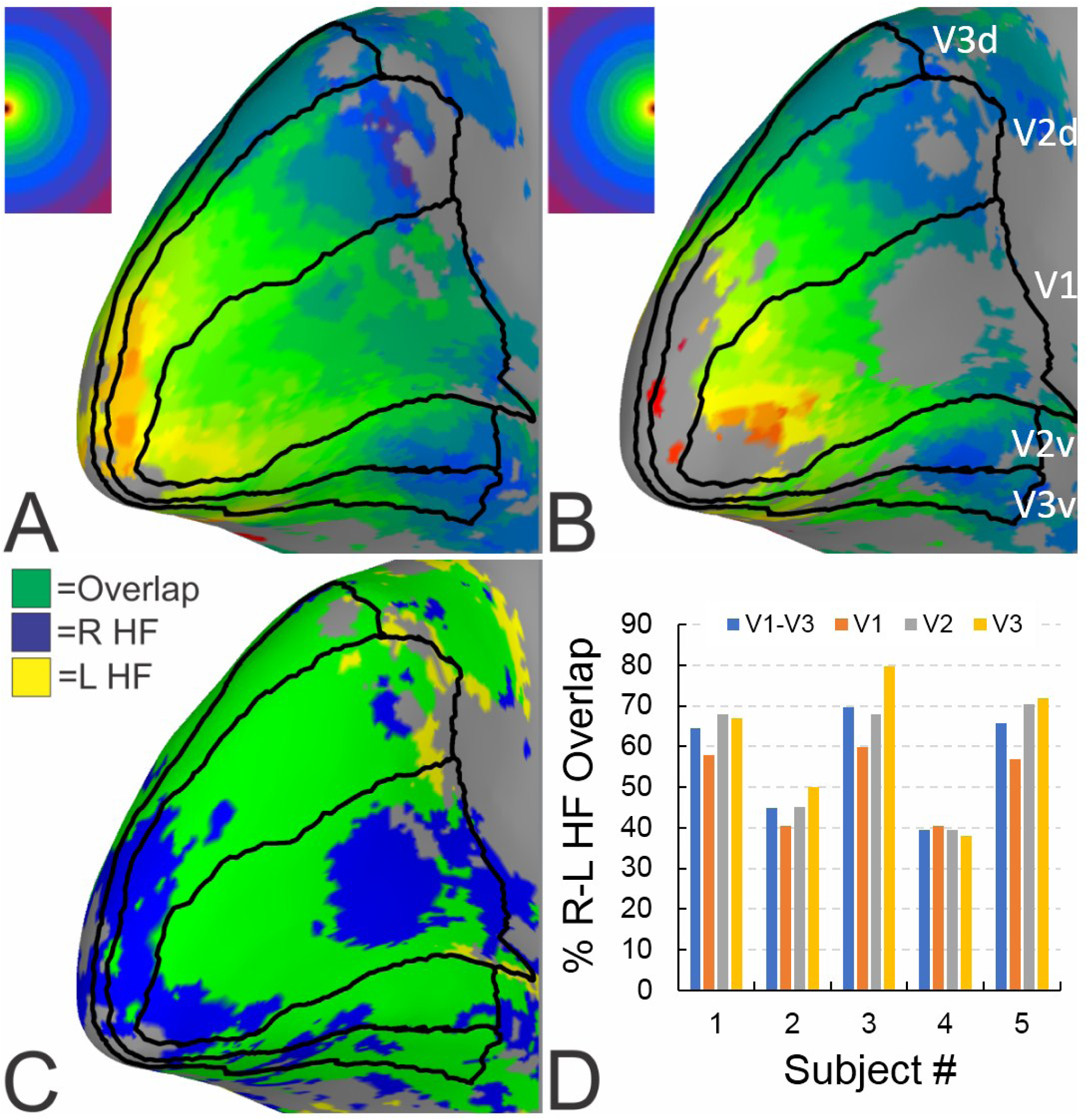
Monocular (right eye) hemifield eccentricity maps and overlap. (A) Right hemifield stimulus condition. (B) Left hemifield stimulus condition. (C) Hemifield overlap map. Green is the logical intersection of A and B after thresholding. (D) Percent hemifield overlap in V1-V3. Data combined across all eye-hemifield stimulus conditions. Color codes for eccentricity and overlap maps are in upper left corner of A-C. Thin black lines demarcate V1-V3 visual area boundaries. All maps are displayed on left occipital cortex surface model for a representative subject with albinism.

The right and left hemifield representations in **Figure 1A,B** were combined logically in **Figure 1C** to form a right-left hemifield overlap map. In this map the blue regions responded exclusively to the right hemifield, yellow regions responded exclusively to the left hemifield, and the green ‘overlap’ regions responded to both hemifields. As displayed in **Figure 1C**, irregular swaths of hemifield overlap (green) are interspersed with right hemifield patches (blue) both ventrally and dorsally in V1, V2 and V3 (black outlines). The extent of hemifield overlap is quantified in **Figure 1D** for V1-V3 in each of our five subjects with albinism. Here, data were combined across all stimuli, eye conditions, and hemispheres. To be classified as “overlapping”, a voxel was required to respond above threshold (*cc* > 0.45) to both a right and left hemifield stimulus regardless of the eye stimulated. Within individual subjects, the percent overlap tended to be similar across V1-V3. However, there were considerable differences in the mean percent overlap across subjects with values ranging from 39%-70%.

### Line of Decussation Shift

To quantify the degree to which the line of decussation shifted in each subject, we estimated the maximum horizontal extent of the aberrant ipsilateral hemifield representations. Using the approach described in (Hoffmann et al., 2003), we drew ROIs for each subject encompassing the horizontal meridian representation along the fundus of the calcarine sulcus in V1. We then made eccentricity histograms for responsive voxels within these ROIs for each of our monocular hemifield stimulation conditions. The maximum eccentricity that encompassed 95% of the data points in the aberrant representation was then used as the measure of decussation shift. To avoid artifactual outliers common at high eccentricities, we limited this analysis to voxels with eccentricity values from 1-15 degrees. These data are listed below in **Table 3**. As the primary aberrant decussation in albinism consists of temporal retinal afferents from each eye decussating to the contralateral hemisphere, we have only listed data in each hemisphere from the contralateral eye stimulation condition. Data for the aberrant ipsilateral field representations in each hemisphere are listed on the left, and the data for the normal contralateral field representations are listed on the right. Aberrant ipsilateral field representations extended horizontally between 3.0-13.8 degrees. This is comparable to previous in reports by Hoffmann et al. in which the aberrant representations extended 5.5-13.9 degrees (Hoffmann et al., 2003). By comparison, the normal contralateral representations extended between 12.5-14.7 degrees. Because we computed the 95% threshold as our measure of maximum horizontal eccentricity, these values do not quite extend to 15 degrees in the normal representations. We were not able to compute this metric in our controls as they did not undergo the hemifield stimulation paradigm; however, Hoffmann et al. estimated that ipsilateral field representations in controls extend horizontally between 0-4.1 degrees eccentricity (Hoffmann et al., 2003).

**Table 3:** Estimation of aberrant decussation extent in albinism

| Subject # | <u>Max Horizontal Ipsilateral Field</u> |  | <u>Max Horizontal Contralateral Field</u> |  |
| --- | --- | --- | --- | --- |
|  | <u>Eccentricity (degrees)</u> |  | <u>Eccentricity (degrees)</u> |  |
|  | Left Hemisphere | Right Hemisphere | Left Hemisphere | Right Hemisphere |
|  | Right Eye<br>Left Hemifield | Left Eye<br>Right Hemifield | Right Eye<br>Right Hemifield | Left Eye<br>Left Hemifield |
| 1 | 9.7 | 13.8 | 14.0 | 13.5 |
| 2 | 4.3 | 3.0 | 13.3 | 14.7 |
| 3 | 6.7 | 7.9 | 13.4 | 13.6 |
| 4 | 6.1 | 3.9 | 14.1 | 13.5 |
| 5 | 9.4 | 8.8 | 12.5 | 13.4 |

### pRFs in Albinism

The partial overlap of opposite hemifield representations in visual cortex gives rise to a complex mix of voxels with either single or dual responses to the rotating wedge stimulus. As illustrated in **Figure 2A** (left), the unusual dual responses (green traces) were easily identified in the rotating wedge timecourse data since they showed ten robust peaks even though the wedge rotation had only five cycles. Yet, other voxels within the same individual responded to the same stimulus with the more typical 5 peaks (**Figure 2A** right). These results are consistent with some voxels responding to two visual field loci per cycle while other voxels respond to a single locus per cycle. To test this quantitatively, we fit each voxel’s time course with both a single Gaussian and a dual Gaussian pRF model. In the top row of **Figure 2A**, we display the time course predicted by the single pRF model (black trace) and in the bottom row we display the dual pRF prediction.

**Figure 2:**
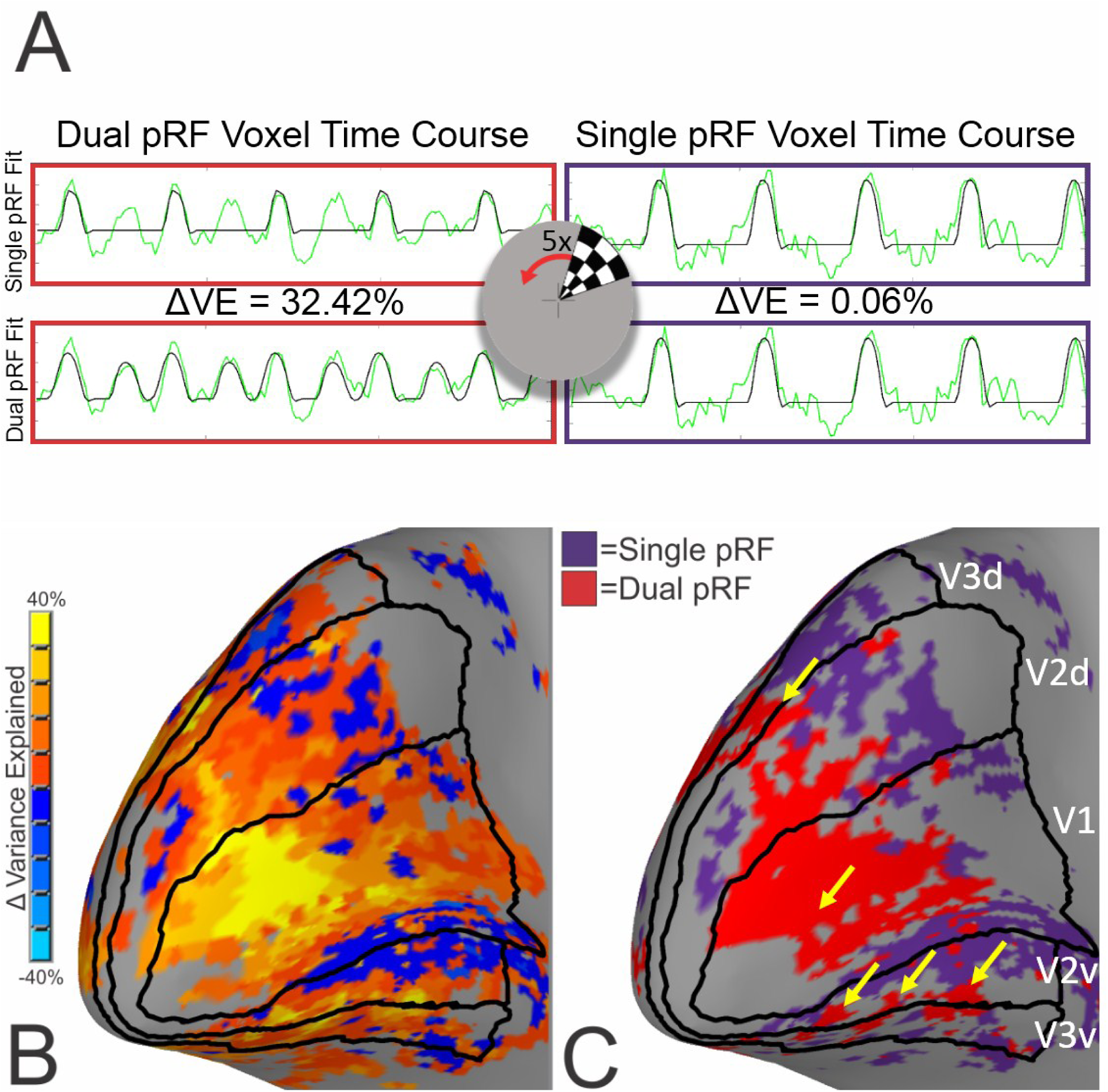
Time courses and distribution of single vs. dual pRF voxels for one subject with albinism. (A) Representative fMRI timecourses (green) for one voxel classified as having a dual pRF (left, red outline) and one voxel classified as having a single pRF (right, purple outline). Black traces represent the single (top) vs. dual (bottom) pRF model predictions and Δ*VE* is the difference in variance explained by the two models (Δ*VE* = *%VE*_*dual*_ - *%VE*_*single*_). Center overlay depicts a single frame of the rotating wedge stimulus. (B) Brain map of ΔVE with color code at left. (C) Distribution of single (purple) vs. dual (red) pRF voxels. Arrows point to dual pRF clusters observed to repeat along horizontal meridian representations.

For the voxel illustrated in the left panel of **Figure 2A**, the dual Gaussian pRF model is able to fit all ten response peaks whereas the single Gaussian model only fits half the peaks. As a result, there is a substantial difference in variance explained (*ΔVE* = 32.42%) by the two models (77.25% vs. 39.83%, *F =* 11.4, *p <* 0.001). However, for the voxel timecourse shown in the right panels, there are only 5 peaks to fit. Consequently, the additional Gaussian parameters of the dual model provide less benefit, and there is little difference in variance explained (*ΔVE* = 0.06%) by the two models (70.04% vs. 69.97%, *F =* 0.02, *p >* 0.999). (Note: Since the two pRF components are fit independently, they tend to be forced into the same visual field locations when the empirical waveform has only five peaks, thus the two models tend to explain the same amount of variance.)

**Figure 2B** shows the voxel-wise difference in variance explained by the two models as a colored pattern across the cortical surface for subject 5. There are clearly large clusters of voxels (yellow patches) in which the dual pRF model explains substantially more variance than the single pRF model, sometimes as much as an additional 40% of the total time course variance for the voxel.

**Figure 2C** displays the results of the partial *F*-test from the comparison of the two models (see Methods) for the same subject. In this analysis, “dual pRF voxels” (red) must have an *F*-value above the 99% confidence interval, and single pRF voxels (purple) must have an *F*-value below the 1% confidence interval. It is notable that dual pRF voxels occurred in large contiguous clusters often, though not always, near the horizontal meridian representations in V1 and on the V2-V3 boundary as indicated by the yellow arrows in **Figure 2C.** In contrast, the single pRF voxels tended to cluster near the vertical meridian representations.

### Incidence and Cortical Distribution of Single and Dual pRF Voxels

Voxels meeting the dual pRF criteria were observed in V1, V2, and V3. **Figure 3A** displays examples of the rotating wedge time courses from several dual pRF voxels in V1-V3 of a representative subject with albinism. Again, the green traces are the empirical rotating wedge time courses, and the black traces are the model predications. The corresponding dual pRF models are represented in visual space in the right column. Note that these voxels all responded to dual locations in opposite hemifields that were approximately mirrored across the vertical meridian. The mean incidence of dual pRF voxels for both albinism and control groups is summarized in **Figure 3B** for V1, V2, and V3 separately as well as combined. Percent dual pRF incidence was compared across groups and visual areas using a two-way ANOVA. This test showed a significant group effect (*p* < 0.001) but no significant visual area effect (*p =* 0. 592) or interaction. Post-hoc tests showed that subjects with albinism

**Figure 3:**
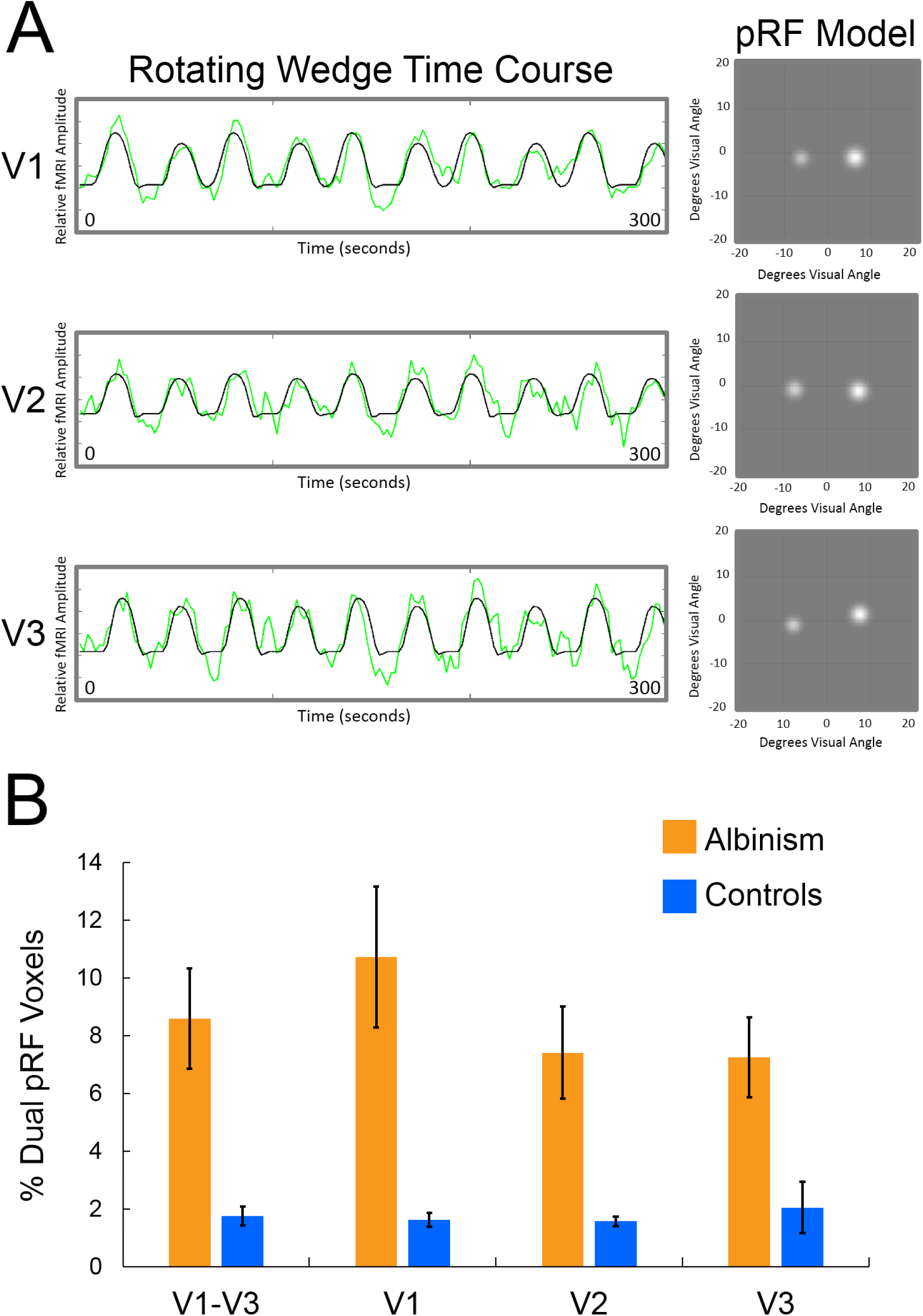
(A) fMRI timecourses (green) and dual pRF model predictions (black) for representative voxels with dual pRFs in V1-V3 of one subject with albinism. Right: Dual pRF models in visual space. White spots represent Gaussian sensitivity profiles for each dual pRF component. (B) Percent incidence of dual pRF voxels in V1-V3 for the albinism and control groups. Subjects with albinism had significantly greater dual pRF incidence in V1, V2, and V3 individually as well as combined across visual areas (respective *p*-values: 0.006, 0.006, 0.013, 0.005). Error bars are *SEM*.

had higher incidences of dual pRF voxels than controls in every visual area. (two tailed, independent t-tests, *p* values for V1, V2, and V3 respectively: *p =* 0.006, *p =* 0.006, *p =* 0.013). On average, dual pRF voxels comprised 8.6% of all visually responsive voxels in the albinism group but ranged from 4% to 15% for different subjects with albinism. This underscores the fact that dual pRF voxel incidence varies across subjects. In contrast, the mean incidence of dual pRFs in control subjects was 1.8% with individual values ranging from of 1% to 3%. Furthermore, visual inspection of many ‘dual pRF’ voxels in controls revealed that they tend to be voxels sampling across the midline or cases in which clear five peaked fMRI responses were overfit by the dual model.

### Dual pRF Symmetry

We assessed the symmetry of dual pRFs by computing the angle formed by the two pRF centers with respect to horizontal. Again, an angle of 0**°** indicates symmetry across the vertical meridian whereas an angle of 90**°** indicates symmetry across the horizontal meridian. These data were sorted into 10**°** bins and histograms were made for each subject plotting the number of dual pRF voxels oriented at each angle. **Figure 4** plots dual pRF angle histograms for the combined group data (top plot) and for each individual subject (bottom plots). Data for subjects with albinism are plotted in orange on the right and control subjects’ data are plotted in blue on the left. The group histograms plot the mean number of dual pRF voxels oriented at each respective angle (error bars = *SEM*), whereas the individual subject histograms plot the total number of dual pRF voxels at each respective angle. As displayed in **Figure 4**, the albinism group’s dual pRF angle data formed a clear peak centered on 0**°.** This pattern is also observable in the data of individual subjects 3, 4, and 5 in the albinism group. The dual pRF angle histogram of subject 1 was more sparsely populated and subject 2 showed a wider distribution of angles indicating a considerable degree of heterogeneity in the range of angles represented in the albinism group. In contrast, the control subjects’ dual pRF angle distributions showed no common central tendency. Dual pRF incidence was compared statistically across both group and angle using a two-way ANOVA. This test showed significant main effects of both group and angle (*p* < 0.001, *p* = 0.002), and a significant interaction between group and angle (*p* = 0.004). The group histograms in **Figure 4** clearly show that the main angle and group effects and the angle*group interaction are driven by the large group differences between bins -30 to 30 with the largest difference occurring at the 0 degree bin. Consequently, we focused all further analyses only on dual pRF voxels with angles between - 30 and 30 degrees.

**Figure 4:**
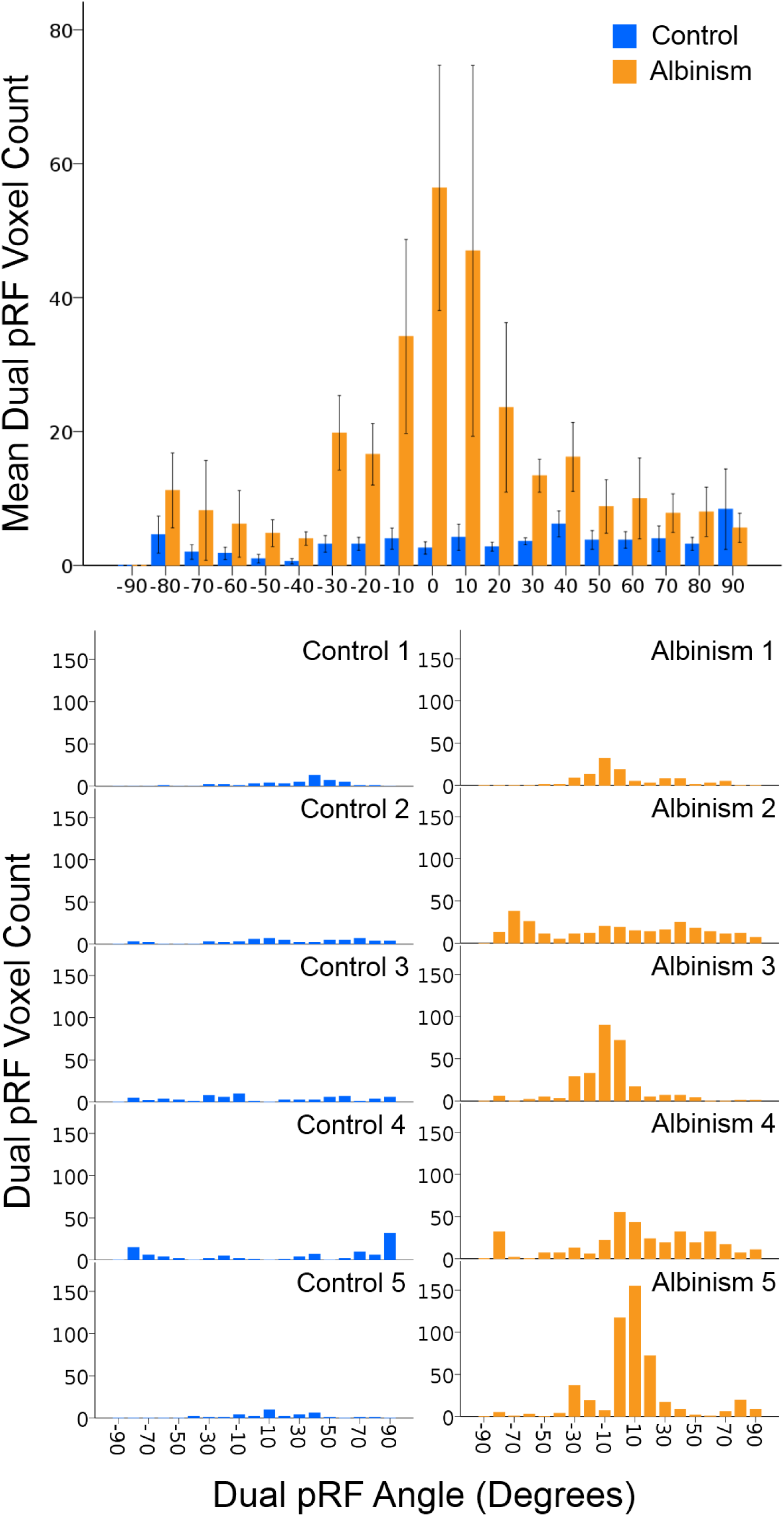
Histograms of dual pRF angle for groups (top) and for individual subjects (bottom). Data for subjects with albinism are plotted in orange on the right, and control subjects’ data are plotted in blue on the left. The top histogram shows the group mean dual pRF voxel count vs. angle formed by the dual pRF centers (degrees). Error bars are *SEM*. Lower histograms show total dual pRF voxel count vs. angle.

### Dual pRF Distribution Relative to Hemifield Overlap Zones in Albinism

To further investigate whether the overlapping right and left hemifield representations in albinism are responsible for aberrant dual pRFs, we examined the association between dual pRF voxels and the cortical zones of R-L hemifield overlap. Note that the overlap zones were computed using the hemifield ring data while the dual pRFs were computed independently using the full field rotating wedge data. **Figure 5A** shows cortical surface models of subject 5’s overlap map (left, repeated from **Figure 1C**) with the corresponding map of single vs. dual pRF voxels (right, repeated from **Figure 2C**). Note that the large red patch of dual pRF voxels in V1 is closely associated with a green patch of hemifield overlap (see arrows). Dual pRF patches on the ventral V2-V3 border are also closely associated with the green hemifield overlap zone in that region. However, note that single pRF zones (purple) also sometimes intersect with the overlap zones indicating that not all voxels in the overlap zone have dual pRFs.

**Figure 5:**
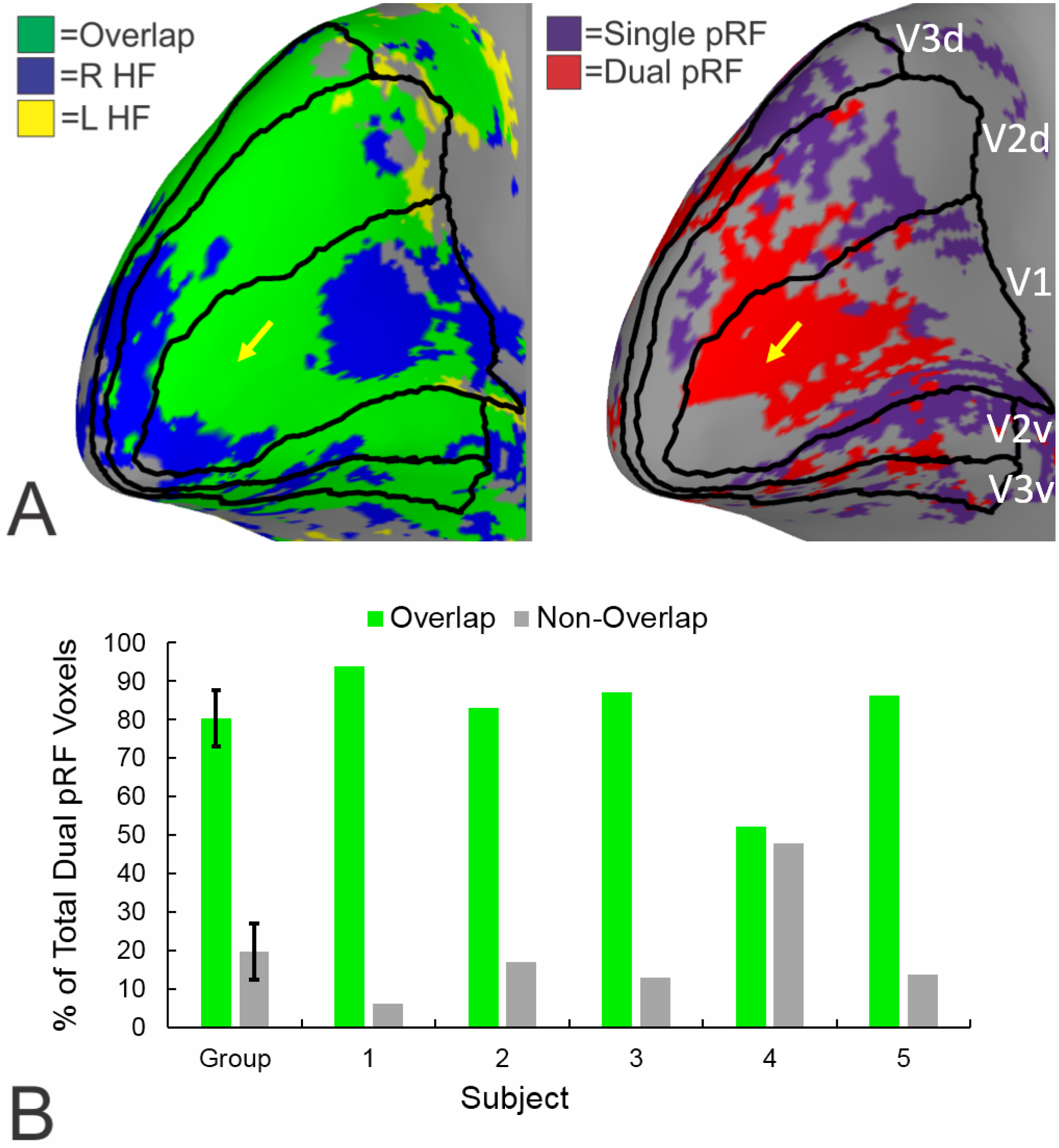
Dual pRF voxels often coincide with right-left hemifield overlap zones. (A) Repeat of Figures 1C and 2C. Note close correspondence of red dual pRF cluster (right arrow) with large green overlap zone (left arrow). (B) Proportion of total dual pRF voxels falling in overlap vs. non-overlap zones for all five subjects with albinism. Error bars are *SEM*.

To quantify the correspondence between the dual pRF clusters and HF overlap zones, we calculated the proportion of dual pRF voxels which fell in non-overlap versus overlap zones (**Figure 5B)**. To be included in this analysis, voxels must have met the inclusion criteria for both the overlap and the pRF analysis described in Methods and have dual pRF angles between -30 and 30 degrees. For the subject whose data are displayed in **Figure 5A** (albinism subject 5), the vast majority (∼86%) of dual pRF voxels fell in the overlap zones. This relationship held strongly for four of the five subjects with albinism: in subjects 1, 2, 3, and 5 the vast majority (83-94%) of dual pRF voxels fell in R-L hemifield overlap zones. However, in subject 4 only 52% of dual pRF voxels fell in the overlap zone. Nonetheless, for the albinism group combined, a significantly greater proportion of dual pRF voxels fell in the hemifield overlap zones than in the non-overlap zones (80% vs. 21%, two sample, two tailed t-test *p =* 0.0004). This was not the case for single pRF voxels (60% vs. 40%, *p =* 0.09).

### Distribution of Dual pRFs Across the Visual Field

To assess the distribution of dual pRFs across the visual field, we plotted each albinism subject’s dual pRFs in visual space in **Figure 6.** We limited these plots to display dual pRFs at eccentricities between 1-10 degrees as 95% of dual pRFs fell within that range. The circle radii represent σ for each dual pRF Gaussian component and dual pRF pairs are connected by a thin line. Note that dual pRFs were not uniformly distributed across the visual field but were instead grouped in idiosyncratic clusters. The total number and extent to which dual pRFs extend outward from the vertical meridian also varies substantially across subjects.

**Figure 6:**
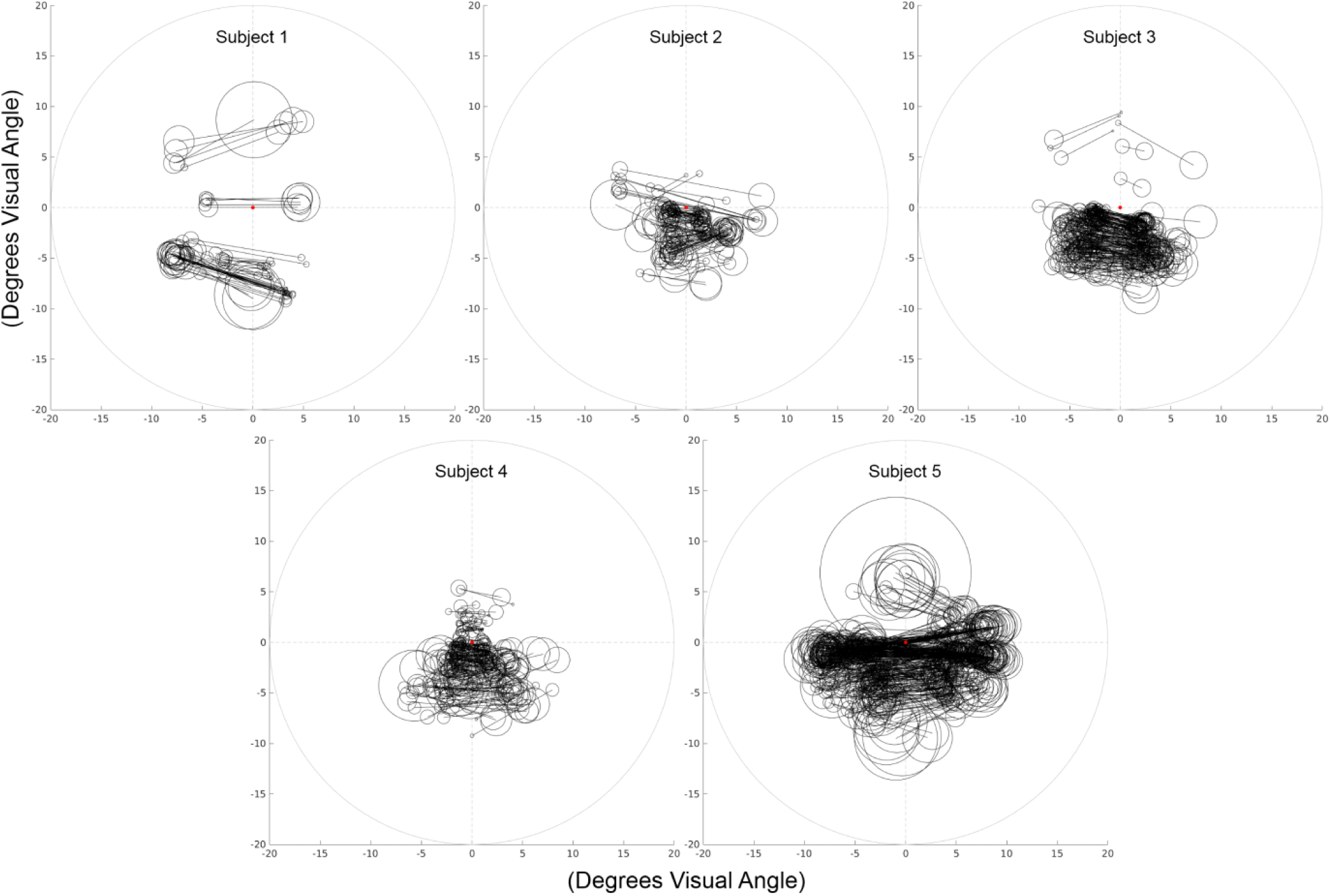
Visual field plots displaying dual pRFs for each subject with albinism. Circle radii represent the sigma for each dual pRF Gaussian component. Thin lines connect each dual pRF pair. Plots display all dual pRFs located between 1-10 degrees eccentricity with angles between -30 and 30 degrees.

### Single and Dual pRF Size Scaling

To measure how single and dual pRF sizes scale with eccentricity, we collapsed each subject’s data into 1 degree eccentricity bins and then plotted mean pRF size (σ) vs. eccentricity for each bin. Data were plotted separately for V1-V3. These data are displayed for V1-V3 of both albinism and control subjects in **Figure 7** (error bars = *SEM*). The first five plots in each row display individual subjects’ pRF size scaling data, and the final plot in each row displays linear trendlines fit to the data pooled across subjects for each group. A color key in the upper left indicates color of symbols representing each visual area.

**Figure 7:**
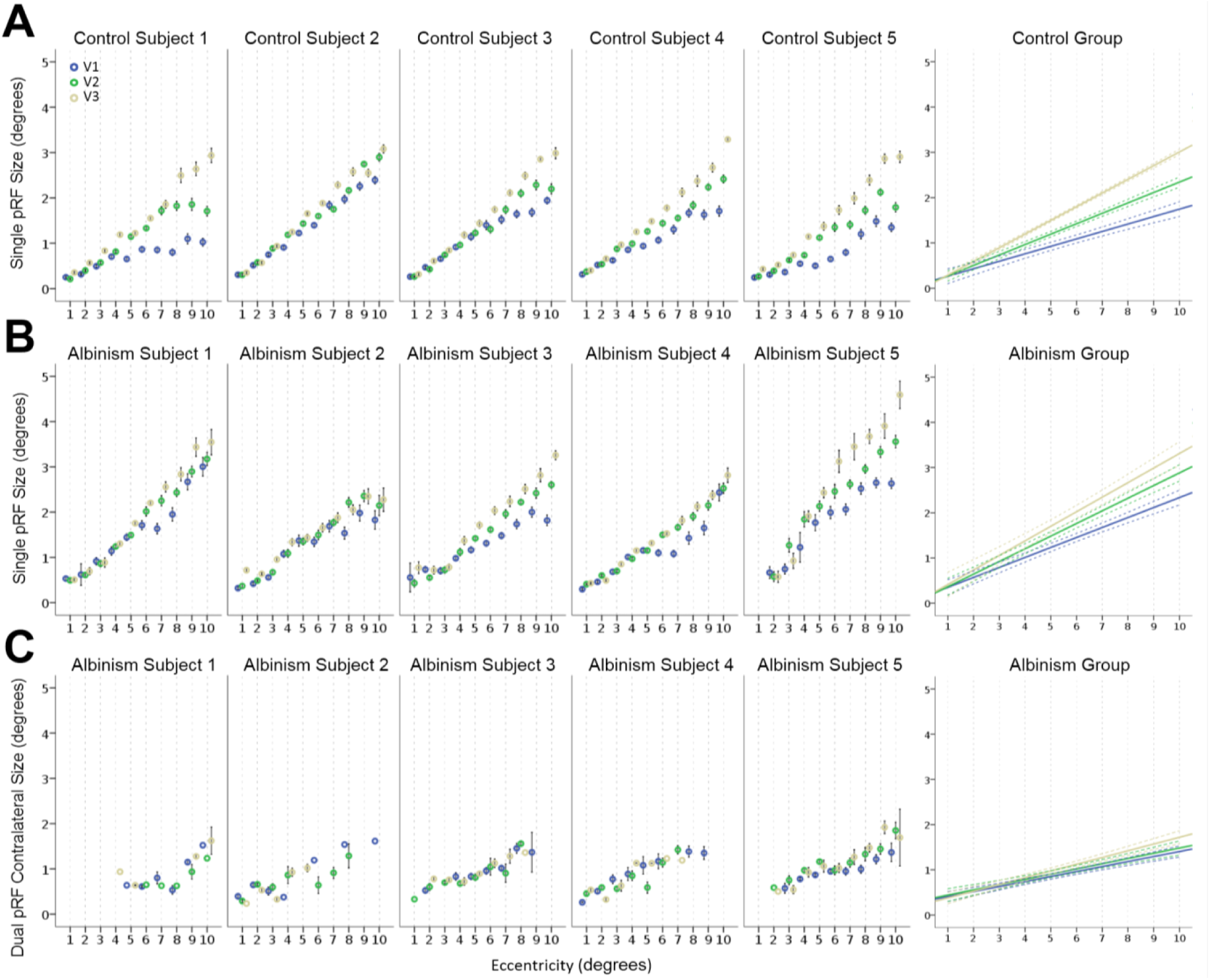
pRF size vs. eccentricity for V1-V3 in albinism and controls. A visual area color code is displayed in the upper left. The first five plots in each row display individual subjects’ pRF size data. The final plot in each row shows linear trend lines fit to the data pooled across subjects. Solid lines are linear trend lines fit for each visual area. Colored dotted lines demark the 95% confidence interval. (A) Single pRFs in control subjects. (B) Single pRFs in subjects with albinism. (C) Contralateral dual pRF component in subjects with albinism. Error bars in individual subject plots are *SEM*.

**Fig 7A** and **B** display single pRF size scaling data for the control and albinism groups respectively, and **Figure 7C** displays dual pRF component size scaling for the albinism group. A three factor, univariate ANOVA (dependent variable: pRF size, factors: component laterality, visual area, and eccentricity) revealed no significant effect of component laterality (ipsi vs. contra) on dual pRF component size across our subjects with albinism (*p =* 0.08). Therefore, we have only displayed the contralateral component sizes. These analyses were also limited to eccentricities of 1-10 degrees for the same reasons stated above. Linear trendlines were fit for each visual area in the group data plots to portray size scaling slopes. Colored dotted lines in the group plots demark the 95% confidence interval of the group mean at each eccentricity.

Single pRF sizes were compared statistically across eccentricity, visual area, and group using a three-factor univariate ANOVA. As expected, there were highly significant main effects of both eccentricity and visual area (*p <* 0.001 for each) and a significant interaction between eccentricity and visual area (*p <* 0.001). These results reflect the consistent scaling of single pRF size with eccentricity and visual area across the control and albinism groups clearly visible in **Figure 7A** and **B.** Subsequent two-factor ANOVAs run on the control and albinism groups separately revealed that the main effects of eccentricity and visual area were significant for both groups individually (all *p* values < 0.001) but that the interaction between eccentricity and visual area was only significant for the control group (*p <* 0.001). Post-hoc Tukey HSD multiple comparisons tests comparing mean single pRF size in each visual area confirmed that the mean pRF size in visual areas V1-V3 were all significantly different from one another in both the albinism and control groups with single pRF sizes increasing with visual area.

Our initial three factor ANOVA comparing single pRF size across eccentricity, visual area, and group also revealed a significant main effect of group (*p <* 0.001) with albinism group having larger single pRFs overall. As displayed in **Figure 8A**, single pRFs were larger in albinism across V1-V3. These mean differences were significant for V1 (1.36 vs. 1.01 degrees) and V2 (1.63 vs. 1.31 degrees) respectively (two tail independent sample t-test, *p =* 0.004 and *p =* 0.034 respectively), but not in V3 (1.87 vs. 1.65 degrees, *p =* 0.215). As illustrated in **Figure 8B**, the group differences for V1 and V2 were also consistent across eccentricity.

**Figure 8:**
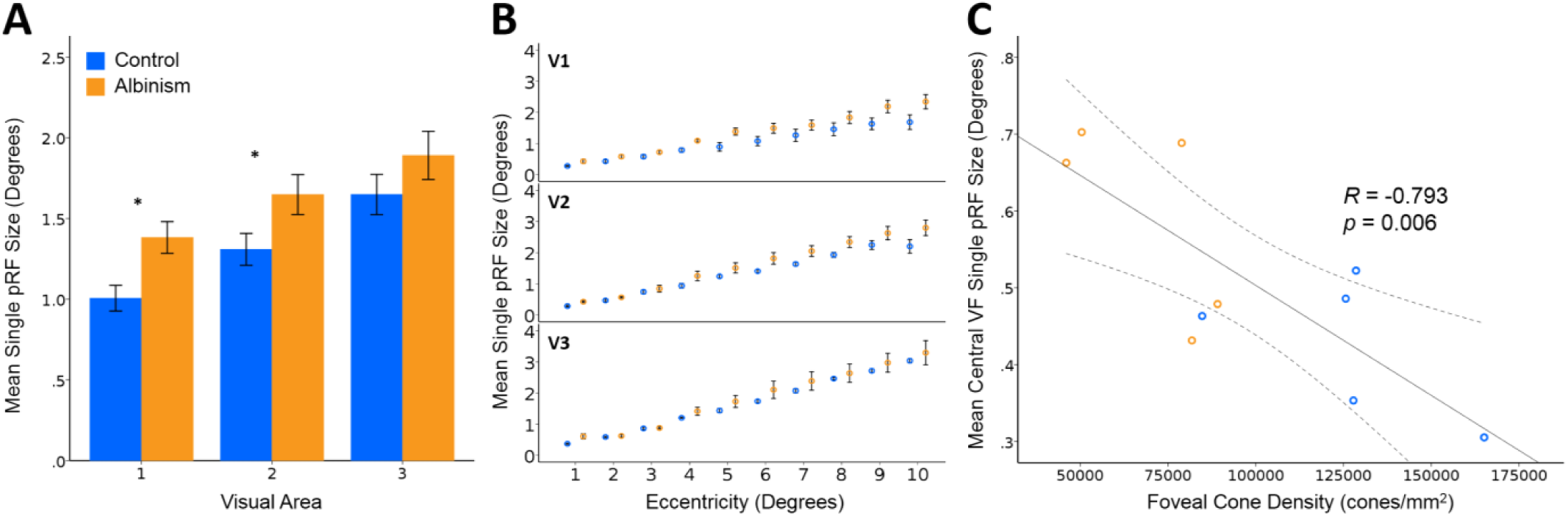
Mean pRF size in the control (blue) and albinism (orange) groups versus visual area (A), eccentricity (B), and foveal cone density (C). Error bars are *SEM*. Dotted lines in (C) represent the 95% confidence interval. (C) displays mean pRF sizes from central visual field (VF) between 1-3 degrees eccentricity for each subject as these locations are nearest to where cone density was measured. The thin black line represents the linear regression fit to the data. *R* is Pearson’s correlation.

To test whether pRF size was related to retinal cone density (Wilk et al., 2014; Wilk et al., 2017; Woertz et al., 2020), we correlated subjects mean pRF sizes between 1-3 degrees eccentricity with their peak foveal cone densities measured by AOSLO in our previously published studies (Wilk et al., 2017; Woertz et al., 2020). We used central field pRF sizes between 1-3 degrees eccentricity as these are closest to where cone density was measured. As displayed in **Figure 8C**, there was a significant negative correlation between foveal cone density and central field pRF size (*R =* -0.793, *p =* 0.006). Whether low cone density in albinism causes larger single pRF sizes in the albinism group is unclear but should be considered in future studies.

Finally, we compared dual pRF component sizes to single pRF sizes in the albinism group across eccentricity and visual area in a univariate three factor ANOVA (Dependent variable: pRF size; Factors: Type (single/dual), eccentricity, visual area). This test again showed significant eccentricity and visual area effects (*p <* 0.001 for each), but also revealed a significant effect of pRF type (*p <* 0.001) and significant interactions of eccentricity*visual area, eccentricity*type, and visual area*type (*p =* 0.028, *p <* 0.001, and *p <* 0.001 respectively). These effects are evident when visually comparing the dual pRF component scaling data to the single pRFs in **Figure 7** as the dual components are consistently smaller than their single pRF counterparts and also scale less steeply with eccentricity and visual area. As mentioned previously, a subsequent ANOVA comparing the size of contralateral and ipsilateral dual pRF components across eccentricity showed no significant effect of component laterality (*p* = 0.08). However, there were, significant main effects of eccentricity (*p <* 0.001), visual area (*p =* 0 .027), and an interaction between eccentricity and laterality (*p =* 0.002). Nonetheless, our post-hoc Tukey test comparing dual pRF component sizes of each visual area failed to show significant differences in dual pRF component size between any pair of visual areas. This is consistent with the lack of dual pRF size differences across visual areas evident in **Figure 7C.**

### Correlation of Dual pRF Voxel Incidence and Fixation Stability

There was also a significant relationship between dual pRF incidence and fixation stability. We correlated dual pRF voxel incidence with subjects’ mean 95% BCEA values for the right and left eye measured in our OPKO fixation stability procedure (**Figure 9)**. Again, this analysis was run only on dual pRF voxels whose pRF angles fell between -30 and 30 degrees. Percent dual pRF voxel incidence correlated significantly with subjects’ mean 95% BCEA measurements (*R* = 0.978, *p =* 0.004).

**Figure 9:**
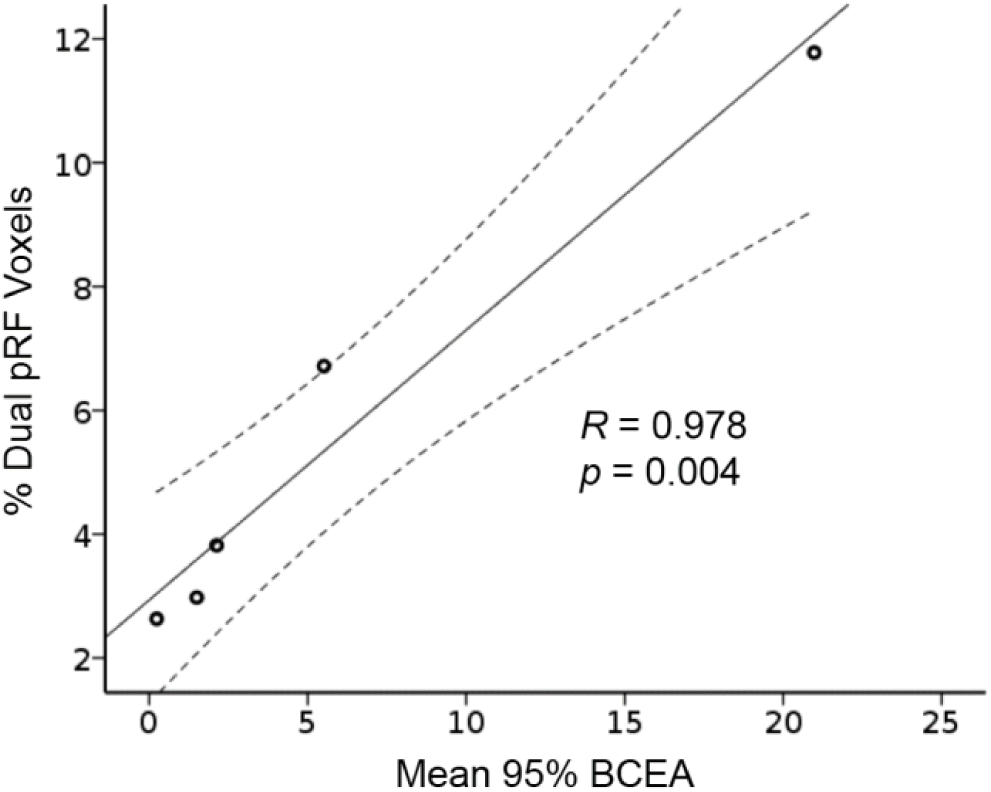
Scatter plot of percent dual pRF incidence vs. mean OPKO 95% BCEA fixation stability for the right eye and left eyes in subjects with albinism. *R* is Pearson’s correlation. The solid line is the linear regression fit. Dotted lines represent the 95% confidence interval.

## DISCUSSION

In this study, we examined the functional organization of visual cortex in subjects with albinism. In particular, we focused on the effects of aberrant retinotopy on characteristics of single voxel pRFs. In all subjects with albinism, we observed partial representations of the ipsilateral visual hemifield that were superimposed on the normal contralateral hemifield representation within each hemisphere, consistent with previous reports (Hoffmann et al., 2003; Kaule et al., 2014; Morland et al., 2001; von dem Hagen, Houston, Hoffmann, & Morland, 2007). We predicted that such overlaid representations would result in single voxels having bilateral dual pRFs. As predicted, we successfully modeled numerous clear cases of dual pRFs in three of our five subjects with albinism (subjects 3, 4, and 5). We also observed many dual pRFs in the other two subjects albeit in smaller, more diffuse clusters. Dual pRF voxels occurred preferentially in cortical zones of R-L hemifield overlap. However, despite the consistent association between dual pRF voxels and the hemifield overlap zones, they were far from coextensive, with dual pRF voxels often comprising a minority of voxels in these regions. To our knowledge, this is the first study to distinguish between dual and single pRF voxels within individual subjects with albinism and report the size scaling of ipsi- and contralateral dual pRF model components. These findings provide clarity in light of recent conflicting reports of whether dual pRF voxels exist in albinism (Ahmadi et al., 2019; Alvarez et al., 2019). In contrast to recent attempts which modeled dual pRFs in albinism explicitly as mirror-symmetrical Gaussian fields, our model optimized the two pRFs independently allowing the two fields to be placed at any pair of angular positions. Even so, our analysis indicated that the two pRF components tend to be positioned approximately (though not always precisely) at mirror image locations across the vertical meridian (**Figure 4**). Our approach of independently optimizing each pRF component was vital to successfully model these not-quite-symmetrical dual pRFs. This approach also allowed us to independently measure size scaling of the contra- and ipsilateral dual pRF components with eccentricity and visual area, and to compare these scaling rates to single pRFs in albinism and controls. Our results indicate that on average single pRFs in albinism are 22% larger than controls, and that the dual pRF components scaled less strongly with eccentricity and visual area than single pRFs in the albinism group. It is also notable that dual pRFs were not uniformly distributed across the visual field **(Figure 6)**. Consequently, tailoring future psychophysical experiments to each albinism subject’s unique dual pRF map may prove useful in detecting potential perceptual consequences of retinocortical miswiring.

### Aberrant Retinotopic Organization in Albinism

We observed substantially overlapping right and left hemifield representations in each cortical hemisphere of all subjects with albinism. This pattern is consistent with the retinal line of decussation (which is normally centered on the vertical meridian) being shifted temporally with the misrouted temporal afferents reaching the same locations in ipsilateral visual cortex as their mirrored nasal counterparts. It’s worth noting that the albinism subtypes represented in our cohort included OA, OCA1B, and OCA2, indicating that such aberrant representations are not unique to any one albinism subtype. The mean percentage of cortex responding to both right and left hemifield stimuli (overlap zones) varied significantly across subjects (**Figure 1**: 39.2-69.7%) as did the line of decussation shift (**Table 3:** 3.0-13.8 degrees). This is consistent with the known variability in extent of miswiring at the optic chiasm across the albinism population as a whole (Creel, Spekreijse, & Reits, 1981; Hoffmann, Lorenz, Morland, & Schmidtborn, 2005; Hoffmann et al., 2003; Kaule et al., 2014; von dem Hagen et al., 2007). The extent of overlap was relatively constant across V1-V3 within individuals (See **Figure 1**) which agrees with previous studies showing that the aberrant ipsilateral representation set up by the retinocortical afferents to V1 is propagated to subsequent stages of the visual hierarchy (Kaule et al., 2014; Wolynski, Kanowski, Meltendorf, Behrens-Baumann, & Hoffmann, 2010).

### Subjects with Albinism Have Voxels with Single and Dual pRFs

Our analysis suggests that the partial superposition of opposite hemifield representations in albinism produces a complex mixture of voxels having either single or dual pRFs. This was qualitatively evident in the rotating wedge time course data as some voxels responded with ten robust peaks (twice per period) while other voxels responded with the expected five peaks (once per period) (See **Figure 2A**). Quantitative modeling indicated that these double responses are best predicted by a dual pRF model, and that dual pRFs are numerous in subjects with albinism (4-15%) but not controls (1-3%). Moreover, a close examination of the pRF fits of such voxels in controls showed that these cases were not convincing, but were instead instances of over fitting and infrequent partial-voluming across the midline. The comparable incidence of dual pRF voxels across V1-V3 in albinism again supports the idea that the aberrant ipsilateral field representation set up by peripheral miswiring is propagated up the visual hierarchy.

### Dual pRF Symmetry

Dual pRF angles in our albinism group peaked at 0 degrees indicating that, the two fields tended to be symmetric across the vertical meridian **(Figure 4)**. These results affirm previous literature that suggests opposite hemifield representations in albinism are superimposed in a mirror symmetrical manner (Hoffmann et al., 2003). However, it is worth noting that some individual dual pRFs are “tilted” by as much as 30 degrees relative to horizontal suggesting that the overlap of the opposite hemifield representations may not always be in perfect retinotopic register, at least within localized regions. This was also noted in (Woertz et al., 2020) where there were sometimes significant mismatches in eccentricity mapping over restricted cortical zones. Consequently, in order to best fit all dual pRF responses in albinism, models should not be rigidly symmetrical, but instead incorporate some flexibility as described here or by using the recently developed ‘micro-probing’ technique described by (Carvalho, Invernizzi, Ahmadi, Hoffmann, Renken, & Cornelissen, 2020). On the other hand, the few “dual pRFs” that were observed in controls showed no clear bias towards any angular orientation. This further supports our interpretation that dual pRFs observed in controls likely arise for artifactual reasons such as model over-fitting.

### Distribution of Dual pRF Voxels Relative to Hemifield Overlap Zones

Given the aberrant superposition of opposite hemifield representations within visual cortex of albinism subjects, one might expect that all voxels within these overlap zones would have dual pRFs whereas voxels outside these zones would all have single pRFs. This was not the case. The vast majority of dual pRF voxels in albinism were found in regions of right-left hemifield overlap (**Figure 5)**, and the spatial patterns of dual pRF clusters often closely corresponded to those of the overlap zones. Nonetheless, single pRF voxels were also numerous within overlap zones in all subjects with albinism, indicating that the occurrence of dual pRFs is not obligatory within hemifield overlap regions. However, it is important to note that the successful modeling of dual pRFs could only occur when there was sufficient separation between the two peaks of each dual response (five dual peak pairs); otherwise, the dual pRF model would perform no better than a “wide” single pRF model. If dual pRFs in albinism are symmetric across the vertical meridian as we and other groups have suggested (**Figure 4**) (Hoffmann et al., 2003), then maximal separation between the two BOLD responses to the rotating wedge stimulus should occur near the horizontal meridian representations, and little to no separation should occur near the vertical meridian representations. This is, exactly, what we observed. We suspect that lack of separation between the dual pRF responses explains why dual pRFs were not more numerous near the vertical meridian. As a result, our measure of dual pRF prevalence is likely to be an under-estimate. Furthermore, it was well documented by Tootell et al. (Tootell, Mendola, Hadjikhani, Liu, & Dale, 1998), and was previously noted by Hoffmann et al. (Hoffmann et al., 2003), that voxels along the vertical meridian representation in controls often respond to both right and left hemifield stimuli due to their receptive fields straddling the meridian boundary. Indeed, a closer analysis of single pRF voxels in the overlap zone revealed a tendency for these cases to be large pRFs near the vertical meridian. In this way, some regions of supposed hemifield overlap with single pRFs may be due to ‘meridian straddling’ rather than aberrant superposition of disparate field locations. Finally, the acquisition of the overlap maps and dual pRF modeling data using two different mapping paradigms (monocular hemifield expanding rings vs. full field binocular rotating wedge) may have contributed to their lack of complete correspondence. Nonetheless, the presence of dual pRF voxels in some regions of hemifield overlap but not others may also indicate spatial heterogeneity in the effects of retinocortical miswiring on local cortical physiology.

### pRF Size

Single pRFs in albinism were, on average, 22% larger than in controls. Moreover, mean pRF size within the central 3 degrees was negatively correlated with foveal cone density across our albinism and control subjects (See **Figs 7** and **8**). This suggests that cone density may contribute to single pRF size. In this way, increased single pRF size is possibly a cortical correlate of the low visual acuity that is known to occur in albinism (Wilk et al., 2014; Wilk et al., 2017; Woertz et al., 2020). However, perplexingly, our previous study did not find strong correlations between cone density and acuity in albinism (Wilk et al., 2014). It is possible that the Snellen acuity measures employed in this study were not accurate enough to capture such effects. Clearly a direct comparison between pRF size, cone density, and acuity in individual albinism subjects is needed to settle this issue.

### Limitations, Potential Artifacts, and Caveats

The presence of dual pRFs in albinism was expected given the previously known superposition of opposite hemifield representations in visual cortex. Nevertheless, one must consider potential artifactual causes of dual pRFs. One potential cause might be that individual imaging voxels encompass retinotopically disparate locations across sulci or across the midline. Though it is possible that some optimally situated voxels might show such effects, the dual pRF voxels we observed in albinism often occurred in large clusters far from the midline and were typically positioned entirely within gray matter. Furthermore, if the dual pRFs we observed were solely an artifact of such partial-voluming, then we would have expected to observe comparable numbers of dual pRFs in our control subjects. This was not the case.

One might also wonder if dual pRFs somehow result from fixational instability or nystagmus. Indeed, our OPKO 95% BCEA measure of fixational stability was positively correlated with the incidence of dual pRFs (**Figure 9**). However, it is extremely difficult to imagine how random instability or even nystagmus could artifactually create a population of dual pRF voxels with components that are mirror symmetric and at a variety of positions relative to the vertical meridian. Furthermore, a recent study (Ahmadi et al., 2019) showed that ‘fixation jittering’ mimicking nystagmus in controls fails to produce voxels which are fit better by a dual-rather than single-pRF model. Previous retinotopic mapping studies in other populations with nystagmus have also shown that moderate nystagmus (50% BCEA of 1-3 degrees) has little or no effect on the fMRI response (Baseler, Brewer, Sharpe, Morland, Jägle, & Wandell, 2002). All but one of our subjects’ BCEAs fell within this range. A more likely explanation for the correlation of BCEA and dual pRF incidence is that both reflect aberrant central visual system organization and, consequently, both tend to increase with the severity of the albinism syndrome.

Unexpectedly, dual pRF sizes were, on average, 38% smaller than single pRF sizes in the albinism group. Scaling of dual pRFs with eccentricity and visual area were also less pronounced than for single pRFs. One potential artifact that might cause such differences would be a failure of the BOLD response to return to baseline during the interval between stimulation of the right and left hemifield components. If this were the case, then the BOLD fluctuations we observed would only reflect the tips of the timecourse peaks, and the width of our models would tend to be smaller than the true pRF widths. However, this effect would then tend to become worse as the angular distance between the dual pRF components became smaller (e.g. for dual pRF components close to the vertical meridian). We explicitly checked for such a correlation but found none. We therefore feel that the reduced size and scaling of dual pRFs relative to single pRFs may be true biological effects. However, additional verification using monocular hemifield stimuli or inclusion of longer rest periods to ensure that responses fully return to baseline may help minimize the potential for this type of confound (Ahmadi et al., 2019; Alvarez et al., 2019).

Albinism is a relatively rare condition, and it is even more uncommon to find people with albinism and fixation stable enough for retinotopic mapping. Consequently, obtaining large numbers of suitable subjects is problematic and is a potential limitation of this study. However, we feel that the results reported here are of sufficient interest to warrant future studies that seek to corroborate these findings in a larger subject cohort.

### Albinism is Heterogeneous both Peripherally and Centrally

Finally, there is a great deal of genetic heterogeneity within the albinism population. Different mutations in the melanin synthesis pathway give rise to different subtypes of albinism as well as heterogeneity in retinal morphology (Kruijt, de Wit, Bergen, Florijn, Schalij-Delfos, & van Genderen, 2018; Lee, Woertz, Visotcky, Wilk, Heitkotter, Linderman, Tarima, Summers, Brooks, Brilliant, Antony, Lujan, & Carroll, 2018; Montoliu et al., 2014; Oetting & King, 1999; Patel, Hayward, Tailor, Nyanhete, Ahlfors, Gabriel, Jannini, Abbou-Rayyah, Henderson, Nischal, Islam, Bitner-Glindzicz, Hurst, Valdivia, Zanolli, Moosajee, Brookes, Papadopoulos, Khaw, Cullup, Jenkins, Dahlmann-Noor, & Sowden, 2019; Prieur & Rebsam, 2017; Simeonov et al., 2013; Wilk et al., 2014). However, specific genetic details for individual patients have not been included in the vast majority of previous imaging studies of albinism. In this study, we intentionally selected albinism subjects to represent a diverse sampling of albinism subtypes (**Table 1**) in order to enhance the probability of observing variation in the central organization of visual pathways. Interestingly we saw disrupted central organization even in subjects with the recently described “tri-allelic” form of albinism (Grønskov, Jespersgaard, Bruun, Harris, Brondum-Nielsen, Andresen, & Rosenberg, 2019; Monferme, Lasseaux, Duncombe-Poulet, Hamel, Defoort-Dhellemmes, Drumare, Zanlonghi, Dollfus, Perdomo, Bonneau, Korobelnik, Plaisant, Michaud, Pennamen, Rooryck-Thambo, Morice-Picard, Paya, & Arveiler, 2019; Norman, O’Gorman, Gibson, Pengelly, Baralle, Ratnayaka, Griffiths, Rose-Zerilli, Ranger, Bunyan, Lee, Page, Newall, Shawkat, Mattocks, Ward, Ennis, & Self, 2017). Though our sample is not nearly large enough to draw strong correlations between genetic, retinal and central factors, we did observe variation across our albinism sample in the extent of hemifield overlap, pRF size, incidence of dual pRFs, and the distribution of dual pRFs across the visual field. We speculate that genetic factors responsible for the range of subtypes in albinism may also underlie the variations in retinocortical organization observed in our dual pRF analysis. If so, this suggests that in a larger sample of subjects with albinism, this approach will prove productive for linking genetic and morphological factors with unique patterns of central miswiring not only in albinism but in a variety of inherited vision disorders (Hoffmann & Dumoulin, 2015). Accounting for individual variability in factors such as dual pRF location and topography in albinism may also allow us to better target future tests for potential perceptual effects of retinocortical miswiring, and ultimately aid in our understanding of the perceptual relevance of retinotopic organization within the visual system.

## CONCLUSIONS

This study shows that retinocortical miswiring in albinism results in single imaging voxels with bilateral dual pRFs. Voxels with dual pRFs are numerous in subjects with albinism but not control subjects and occur selectively (but not ubiquitously) in cortical regions where the right and left hemifield representations are superimposed. Our results agree with previous studies which predict that dual pRFs in albinism are positioned at approximately mirror image locations across the vertical meridian but suggest that this symmetry is not always precise. Thus, in order to accurately fit responses from all dual pRF voxels, models cannot be rigidly mirror symmetrical, but must instead incorporate some flexibility.

Our results also show that single pRFs are larger in albinism than in controls, and that subjects’ mean pRF sizes in the central visual field are highly anti-correlated with their foveal cone densities. This suggests that low foveal cone density in albinism may engender larger single pRFs in visual cortex which, in turn, may contribute to the reduced central acuity typical of albinism.

Finally, dual pRFs in albinism were not evenly distributed across the visual field, but instead occurred in idiosyncratic clusters unique to each subject. In the future, mapping subjects’ unique dual pRF distributions could guide spatially focused psychophysical tests aimed at revealing previously undetected perceptual consequences of retinocortical miswiring in albinism.

## ACKNOWLEGEMENTS

The authors would like to thank Melissa Wilk and Erin Curran for their contributions to this work. We would also like to thank Murray Brilliant, Robert Valenzuela, and Jeff Joyce for their contributions to our genetic testing. Research reported in this publication was supported by the National Eye Institute, the National Institute of General Medical Sciences, and the National Center for Advancing Translational Sciences of the National Institutes of Health under award numbers TL1TR001437, T32GM080202, T32EY014537, P30EY001931, and R01EY024969. This investigation was conducted in a facility constructed with support from Research Facilities Improvement Program, Grant Number C06RR016511, from the National Center for Research Resources, National Institutes of Health. The content is solely the responsibility of the authors and does not necessarily represent the official views of the National Institutes of Health. This work was also supported by Vision for Tomorrow.

